# *De novo* assembly of two Swedish genomes reveals missing segments from the human GRCh38 reference and improves variant calling of population-scale sequencing data

**DOI:** 10.1101/267062

**Authors:** Adam Ameur, Huiwen Che, Marcel Martin, Ignas Bunikis, Johan Dahlberg, Ida Höijer, Susana Häggqvist, Francesco Vezzi, Jessica Nordlund, Pall Olason, Lars Feuk, Ulf Gyllensten

## Abstract

We have performed *de novo* assembly of two Swedish genomes using long-read sequencing and optical mapping, resulting in total assembly sizes of nearly 3 Gb and hybrid scaffold N50 values of over 45 Mb. A further analysis revealed over 10 Mb of sequences absent from the human GRCh38 reference in each individual. Around 6 Mb of these novel sequences (NS) are shared with a Chinese personal genome. The NS are highly repetitive, have elevated GC-content and are primarily located in centromeric or telomeric regions. A BLAST search showed that 31% of the NS are different from any sequences deposited in nucleotide databases. The remaining NS correspond to human (62%) or primate (6%) nucleotide entries, while 1% of hits show the highest similarity to other species, including mouse and a few different classes of parasitic worms. Up to 1 Mb of NS can be assigned to chromosome Y, and large segments are missing from GRCh38 also at chromosomes 14, 17 and 21. Inclusion of these novel sequences into the GRCh38 reference radically improves the alignment and variant calling of whole-genome sequencing data at several genomic loci. Through a re-analysis of 200 samples from a Swedish population-scale sequencing project, we obtained over 75,000 putative novel SNVs per individual when using a custom version of GRCh38 extended with 17.3 Mb of NS. In addition, about 10,000 false positive SNV calls per individual were removed from the GRCh38 autosomes and sex chromosomes in the re-analysis, with some of them located in protein coding regions.

## Introduction

Due to advances in DNA sequencing technologies, whole genome sequencing (WGS) has become an established method to study human genetic variation at a population scale. Large human WGS projects have been initiated in several countries and geographic regions^1–6^, in some cases comprising 10,000 individuals or more^7,8^. These genome projects will provide a wealth of information for future research on human genetics, evolution and disease. Today, the vast majority of human WGS is performed using short-read Illumina sequencing technology, and requires an alignment of the sequence reads to a human reference sequence. The gold standard reference is the GRCh38 release from 2013, which is based on DNA from multiple donors and intended to represent a pan-human genome, rather than a single individual or population group^9^. However, the current GRCh38 reference might not be optimal in the context of population specific WGS projects, and more information could be gained from WGS data by instead using local references genomes, tailored to a specific country or population. For instance, the *de novo* assembly of 150 Danish individuals based on Illumina mate-pair sequencing have strengthened the hypothesis that regional reference genomes can increase the power of association studies and improve precision medicine^10^. Since Illumina’s technology is limited by short read lengths and amplification biases^11^ it is not a viable alternative for creating human *de novo* assemblies comparable to GRCh38 in terms of completeness and contiguity.

A number of sequencing technologies have emerged that are capable of reading very long DNA molecules without prior amplification. These methods can resolve complex regions of the genome, such as GC-rich regions or repeats, which are difficult to determine with amplification-based and short-read approaches. In particular, PacBio’s single-molecule real-time (SMRT) sequencing technology has proven to be an excellent method for *de novo* genome assembly. In 2015, the first human *de novo* SMRT sequencing project was reported; the assembly of the CHM1 cell line derived from a haploid hydatidiform mole^12^. Since then, a handful of human genomes have been assembled using combinations of long-read, linked-read and optical mapping technologies^13–16^, including the AK1 cell line originating from a Korean individual^15^ and the HX1 genome originating from a healthy male Han Chinese^16^. These personal genomes have been assembled to high level of completeness. For example, the AK1 assembly has a contig N50 size of 17.9 Mb and scaffold N50 size of 44.8 Mb, with 8 chromosome arms resolved into single scaffolds^15^. The contigs assembled from SMRT sequence data, as opposed to most assemblies based on Illumina data, are completely gap-free contain no unambiguous bases (represented by N’s). Approximately 20,000 structural variations (SVs) are detected by SMRT sequencing of a human individual^15,16^ and a majority of these SVs are missed by analyses of short-read Illumina WGS data^14^. The assemblies generated using SMRT sequencing have also indicated that a substantial amount of sequence is missing from the GRCh38 version of the human reference. For example, 12.8 megabases (Mb) of novel sequences (NS) were detected in the Chinese HX1 assembly^16^. Also, an average of 0.7 Mb per individual not present in GRCh38 were found among the 10,000 samples in a population-scale Illumina WGS project^8^, showing that NS can to some extent be detected also in short read data.

The human *de novo* assemblies available based on long-read data thus indicate that each personal genome contains a significant amount of “dark matter” of structural variation that is not detected by short-read WGS, and several million bases of novel sequence that cannot be matched to GRCh38. At present, it is unknown how many of these SVs and NS are common to all humans, and thus represent errors in the GRCh38 reference, and how many of them are polymorphic between individuals. To address this question there is a need to assemble several personal genomes from different populations around the world to a high degree of completeness. Such a collection of *de novo* genomes would make it possible to improve on GRCh38, and eventually to construct complete new population-specific genomes. In this study, we performed *de novo* assembly of genomes from two individuals from the Swedish population, in order to investigate the missing pieces of GRCh38, and to evaluate the benefits using a local reference for single nucleotide variant (SNV) calling in population-based WGS data.

## Results

### De novo assembly of two Swedish individuals

To construct two high quality genome references for the Swedish population, DNA was extracted from blood samples obtained from one male (Swe1) and one female (Swe2). The two individuals were unrelated and selected from the 1000 samples included in SweGen, which is a project where the genetic variation in a cross-section of the Swedish population was studied using Illumina WGS^5^. A principal component analysis (PCA) shows that Swe1 and Swe2 are relatively distant from each other in the context of the genetic variation within Sweden (**Figure 1A**). The two selected genomes should therefore contain a large portion of the common genetic variation in the Swedish population.

**Figure 1.**
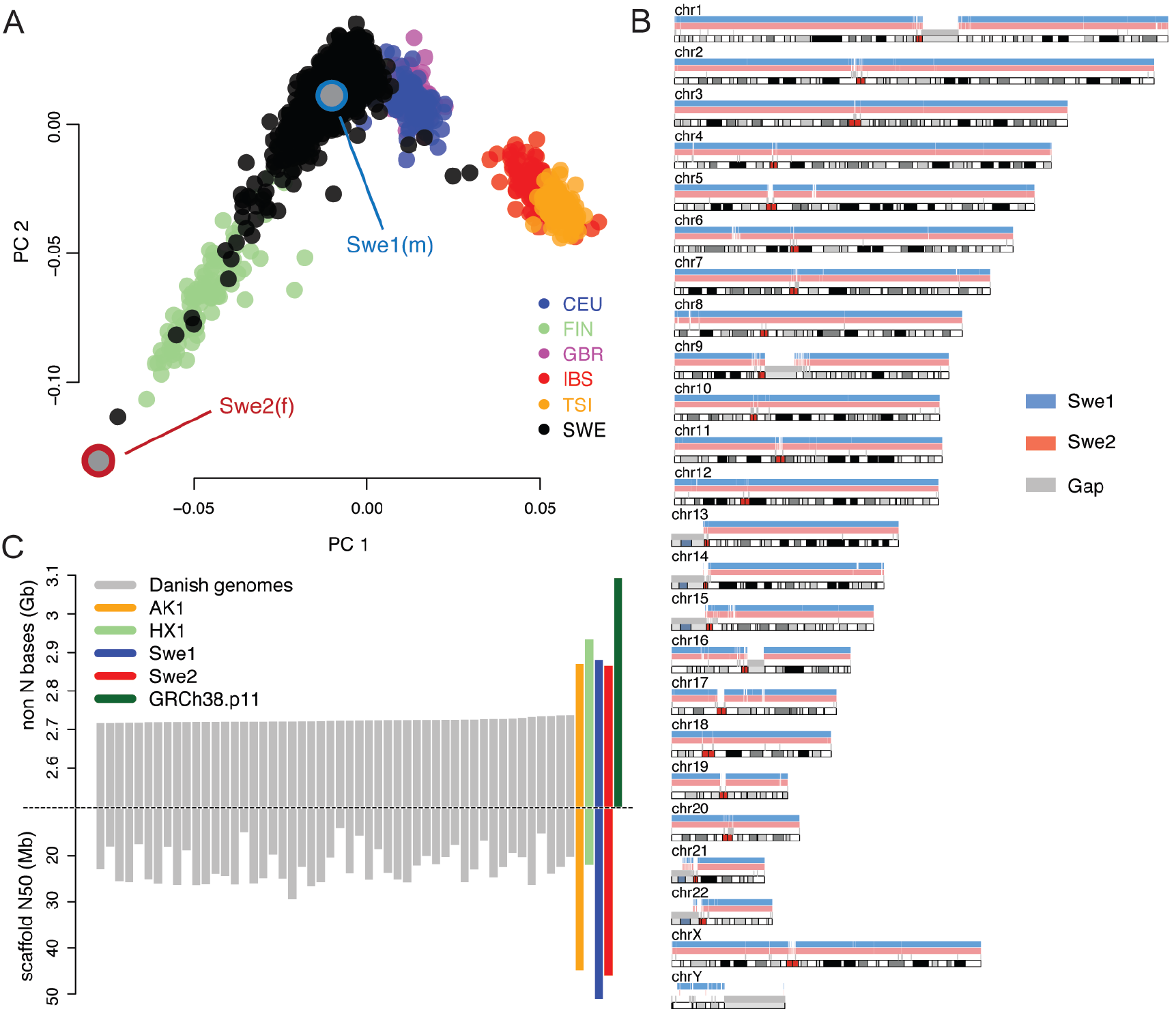
Selection of individuals and *de novo* assembly results. **A)** Results of PCA of WGS data from the SweGen project^5^, compared to the European 1000 Genomes data^32^ (CEU: Utah Residents with Northern and Western Ancestry, FIN: Finnish in Finland, GBR: British in England and Scotland, IBS: Iberian Population in Spain, TSI: Toscani in Italia). The black dots indicate 942 samples from the Swedish Twin Registry (STR), which were sequenced within the SweGen project and represent a cross-section of the Swedish population. Swe1 and Swe2 are the individuals selected for *de novo* sequencing. **B)** Alignment of contigs for Swe1 (blue) and Swe2 (red) to the human GRCh38 reference. 6,812 contigs could be aligned for Swe1 and 6,924 for Swe2. Only the male Swe1 sample has extensive coverage of the Y chromosome. **C)** The bars show the total number of non-N bases (top) and scaffold N50 values (bottom) for Swe1, Swe2 and a selection of other human *de novo* assemblies. The grey bars represent the top 50 genomes with highest number of non-N bases from an Illumina mate-pair assembly of 150 individuals^10^. The Korean (AK1) and Chinese (HX1) genomes were assembled by a combination of SMRT sequencing and optical mapping. Scaffold N50 is not shown for GRCh38 (in green) since it is much higher than for the personal genomes and difficult to fit into the same plot.

SMRT sequencing data was generated at an average coverage of 78.7X for Swe1 and 77.8X for Swe2 (**supplementary Table S1**). By *de novo* assembly^17^, followed by two iterations of genome polishing, we were able to construct sequence assemblies of 2.996 Gb and 2.978 Gb for Swe1 and Swe2, consisting of 7,166 and 7,186 contigs, respectively (**supplementary Table S2**). Each of the assemblies contained about 3,000 primary contigs and an additional 4,000 alternative contigs originating from regions with high heterozygosity. The alternative contigs only cover a small fraction of the genome; about 115 Mb in each individual. N50 values for the primary contigs were 9.5 Mb for Swe1 and 8.5 Mb for Swe2. For both individuals, we also generated BioNano optical mapping data with two different labeling enzymes, at over 100X coverage per enzyme. A two-step hybrid scaffolding of the SMRT sequencing contigs together with the optical maps resulted in assemblies of size 3.1 Gb and scaffold N50 of 49.8 Mb (Swe1) and 45.4 Mb (Swe2) (**supplementary Table S3**). These numbers are similar to the 44.8 Mb scaffold N50 obtained for the first published Korean genome^15^, and substantially larger than the median scaffold N50 of 21 Mb obtained for 150 Danish genomes^10^. It is worth to note that the DNA samples used for optical mapping of Swe1 and Swe2 were extracted from blood collected in 2006. Our results thus demonstrate that it is possible to obtain very high-quality genome assemblies starting from frozen blood that has been stored in the freezer for over a decade.

### Evaluating the quality of the de novo assemblies

To assess the quality of the two *de novo* assemblies, we aligned the contigs for Swe1 and Swe2 to the hg38 reference genome. Throughout this article, the abbreviation hg38 is used to denote a sequence that is identical to the full analysis set of GRCh38, which includes un-localized scaffolds and decoy sequences^18^ (see Methods). For Swe1 and Swe2 respectively, 2.971 Gb (99.14%) and 2.956 Gb (99.24%) of assembled sequence could be uniquely aligned to hg38 (see **Figure 1B** and **Supplementary Table S4**). The slightly higher number of aligned bases for Swe1, who is a male, can be explained by sequences on the Y chromosome that are not present in the female Swe2 sample. The average identity between the contigs and hg38 was over 99.7% for both genomes. Intriguingly, a higher fraction of the Swe2 sequence data can be uniquely aligned to the Swe1 *de novo* assembly (99.55%) as compared to hg38 (99.24%), thus suggesting that the hg38 reference does not contain all sequences present in these Swedish individuals. The corresponding analysis for Swe1 is not relevant in this context, since Swe1 is expected to contain sequence on the Y chromosome not present in the female Swe2.

In order to discover novel sequence (NS) missing from hg38, it is essential that Swe1 and Swe2 were assembled to a high degree of completeness and with as few gaps as possible. To evaluate this, we compared our Swedish *de novo* assemblies to results obtained for the Korean AK1^15^, the Chinese HX1^16^ and 150 Danish genomes^10^. As seen in **Figure 1C**, the primary contigs of Swe1 and Swe2 contain similar number of unambiguous (non-N) bases as the other SMRT sequencing assemblies (i.e. AK1 and HX1). Importantly, the assemblies obtained from SMRT sequencing contain over 100 Mb of additional sequence as compared to the Illumina mate-pair assemblies. The GRCh38 reference contains almost 3.1 Gb, which is significantly more as compared to the ~2.9 Gb for Swe1 and Swe2. To a certain extent, these differences can be explained by the fact that GRCh38 is based on a combination of DNA sequences and haplotypes from several individuals^9^, which could lead to an inflated genome size, and also that primary contigs shorter than 20kb were excluded from the Swe1 and Swe2 assemblies. N50 scaffold values are highest for the Swe1, Swe2 and AK1 assemblies, which all used BioNano data from two labeling enzymes for hybrid scaffolding. A single enzyme was used for HX1 and this assembly has a scaffold N50 similar to those obtained for the Danish genomes.

### Structural variation in Swedish genomes

Analysis of structural variation (SV) resulted in a total of 17,936 SVs for Swe1 and 17,687 SVs for Swe2 (**supplementary Table S5**). These numbers can be compared with the 20,175 and 18,210 SVs detected in the Chinese HX1 and in the Korean AK1 assembly, respectively. The SV length distribution shows an enrichment of ALU repeat elements at around 300 bp and of LINE elements at around 6100 bp, similar to what has been previously reported^15^ (**supplementary Figure S1**).

### Detection of novel sequences not present in the human reference

Even though most of the contigs in our assemblies were in good agreement with the human reference, we detected 25.6 Mb of sequences in Swe1 and 22.6 Mb in Swe2 that could not be aligned to hg38 (**supplementary Table S4**). To refine these sequences further, we performed a two-step re-alignment using more relaxed settings and removed duplicated sequences (see Methods). This resulted in 2859 NS in Swe1 of total length 13.8 Mb, and 2786 NS in Swe2 of total length 10.6 Mb (**supplementary Table S6**). The NS were required to be at least 100bp in length and have at most 80% identity to hg38. They could either originate from contigs that could not be aligned to hg38, or from inserted elements in the aligned contigs. As seen in **Figure 2A**, most of the NS are relatively short (between 100bp and 5kb). However, 83% of NS bases originate from sequences that are over 5kb in length.

**Figure 2.**
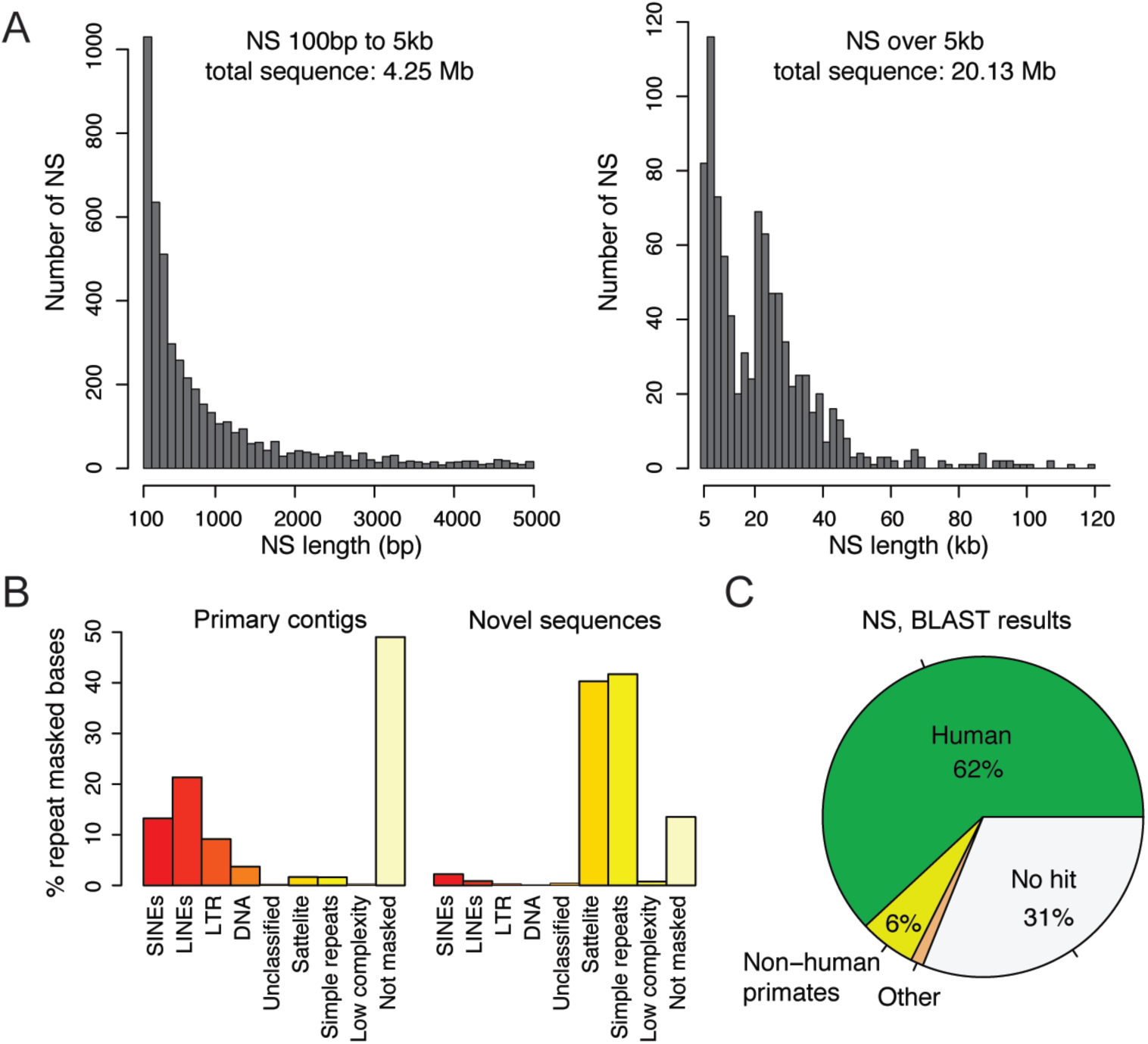
Characterization of novel sequences found in Swe1 and Swe2. **A)** The histograms show the length distribution of all NS found in Swe1 and Swe2. Shorter NS are displayed in the left panel (100bp to 5kb), and longer NS are shown in the right panel (>5 kb). The longer NS comprise the majority of the NS in Swe1 and Swe2. **B)** Results of repeat masking in primary contigs (left) and NS (right). Within the primary contigs 51% of the bases are found to be repetitive using the repeat masker software, while 86% of the bases are repetitive within the NS. Satellite and simple repeats makes up 82% of the bases in the NS. **C)** Results of matching the 5,645 NS in Swe1 and Swe2 to the NCBI database using BLAST^28^. Each piece of the pie chart represents the number of NS that were assigned to a particular species as the top hit. The ‘No hit’ category (in white) contains NS where no E-value reached 1e-50 or lower. 72 of the NS are in the ‘Other’ category, which includes matches to a number of parasitic worms (both for Swe1 and Swe2) and a complete HPV35 genome (only for Swe2).

Repeat masking using sensitive settings, revealed an abundance of repetitive elements in the NS (see **Figure 2B** and **supplementary Table S7**). For Swe1, 88.58% of the NS bases were found to be repetitive. A slightly lower repeat content, 83.60% was detected in the NS from Swe2. Since the repeat content is around 50% among all Swe1 and Swe2 contigs, there is a high enrichment of repeats in the NS. Also, the GC level is slightly elevated, 42.68% (Swe1 NS) and 43.45% (Swe2 NS), as compared to 40.95% in all contigs (**supplementary Table S8**). SMRT sequencing is known to perform well in repetitive regions and high-GC regions, and therefore these results are not unexpected. Annotation of all the repeats showed that satellites and simple repeats make up 82% of the NS bases, but only 3% of all of the primary contig sequences (see **supplementary Table S7**). Interestingly, all other groups of repeats are underrepresented among the NS. Even though a high proportion of the NS are repetitive, there is also a substantial amount of non-repetitive sequence. For Swe1 and Swe2, 1.58 Mb and 1.73 Mb of NS remained after the repeat masking.

### Origin of the novel sequences

To further investigate the contents of the novel sequences, we performed a BLAST search^19^ against all sequences present in the NCBI database (see **Figure 2C**, **supplementary Table S9**). Thirty-one percent of the NS did not produce a BLAST hit, implying that they have not been previously reported. For the remaining NS, the majority matched to human entries in NCBI, thereby suggesting that a majority of our NS have been detected in previously, but originate from regions or haplotypes that have not been included in the hg38 reference. We also detected 5% non-human primate sequences, which most likely originate from regions missing in hg38 that have been sequenced in another primate. Nearly 1% of the NS match to other, non-primate, species. Of note, several of the hits show high similarity to parasitic worms, including *Spirometra erinaceieuropaei*, *Enterobius vermicularis* and *Dracunculus medinensis*. Since it is highly unlikely that the two Swedish individuals indeed have DNA from these parasites present in their blood, a more plausible explanation is that the worm genome assemblies contain a fraction of human sequence. The initial worm assemblies were based on short-read sequencing of samples extracted from human patients^20^, and contigs not aligning to GRCh38 may have been mis-annotated as worm sequence, thus explaining the overlap with our NS. Notably, the Swe2 sample also contained a complete human papilloma virus 35 (HPV35). This could either originate from an HPV35 infection in the blood of this individual, or from a contamination in the sample.

### Comparing novel sequences between Swedish individuals and the Chinese HX1

To investigate whether the NS are individual-specific, population-specific or shared between different populations, we compared our results to those obtained for the Chinese HX1 genome^16^. The HX1 assembly was based on 103X genome-wide SMRT sequencing of DNA from a human blood sample, where 12.8 Mb of NS was found. Starting from the NS identified in Swe1 or Swe2, we determined whether the same sequences could be identified in the other Swedish assembly, or in HX1 (see **Figure 3A** and **supplementary Table S10)**. For Swe1 and Swe2, 55% (7.65 Mb) and 52% (5.51 Mb) of the NS, respectively, could also be found in the other Swedish individual as well as in HX1. The higher overlap obtained when starting the analysis from Swe1 is explained by certain repetitive elements that occur with higher copy number in Swe1 as compared to Swe2. Our results also show the presence of over 5 Mb of NS in the 3-way overlap category, i.e. found in all three individuals. A smaller amount of NS (~1.5 Mb) was common only between the two Swedish individuals, while not found in the Chinese HX1, thus representing possible population-specific sequence. We also identified a substantial amount of individual-specific sequences, 3.27 Mb for Swe1 and 3.22 Mb for Swe2. Interestingly, a much higher amount of NS were shared between Swe1 and HX1 (1.36 Mb) as compared to Swe2 and HX1 (0.29 Mb). Since Swe1 and HX1 are both males, while Swe2 is a female, the ~1 Mb of additional NS shared between Swe1 and HX1 may be explained by segments of the Y chromosome that are missing from the hg38 reference.

**Figure 3.**
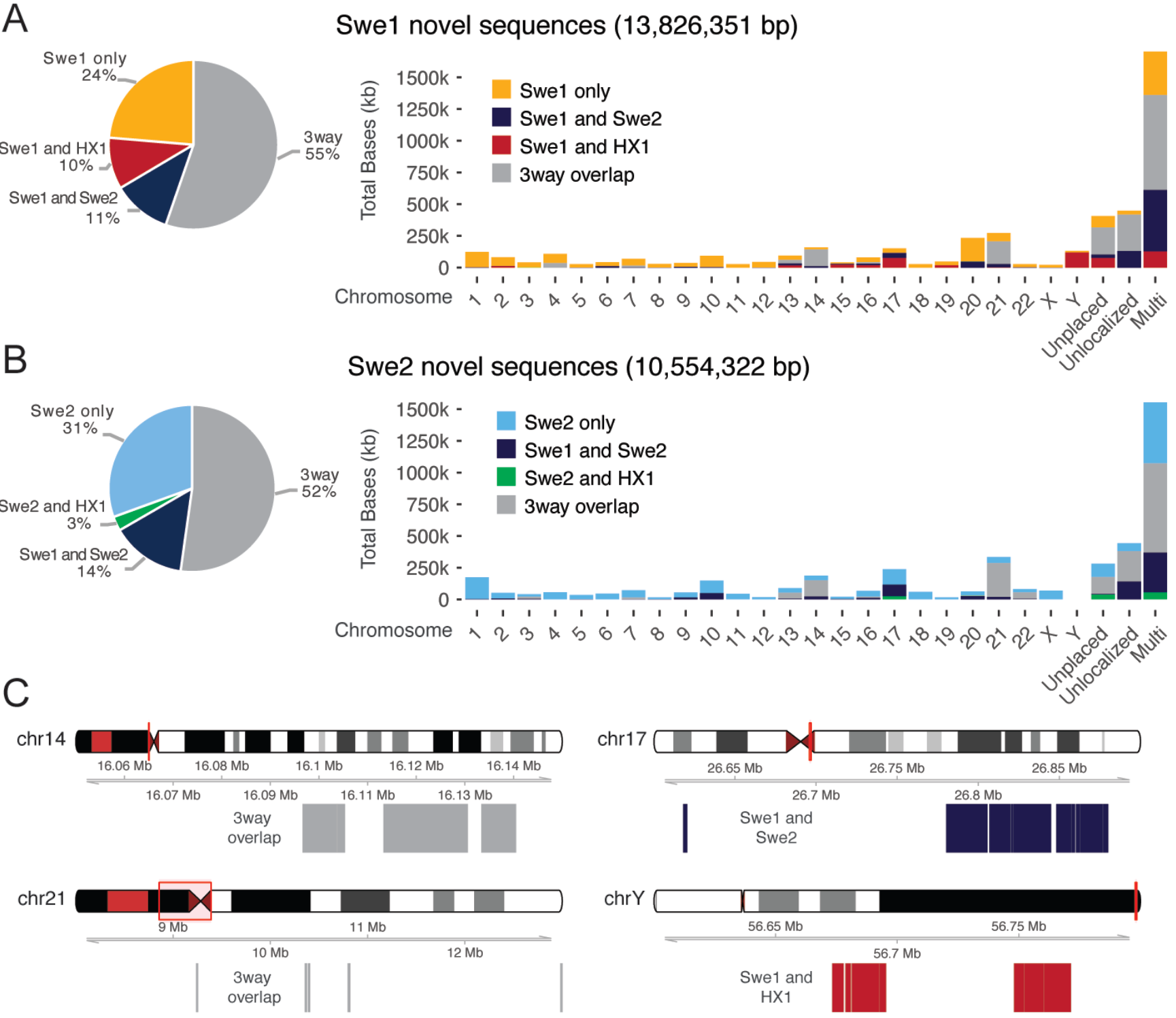
Anchoring of Swe1 and Swe2 NS to the hg38 reference. **A)** The pie chart to the left shows the proportion of Swe1 NS (in total 13,8 Mb) that are also found in Swe2 or in the Chinese HX1. The category 3way (grey) represents NS that are found in all three individuals. The bars to the right shows the amount of NS that can be anchored to the hg38 genome. The category ‘unplaced’ represents sequences in hg38 that are not associated with any chromosome, and ‘unlocalized’ corresponds to sequences that are associated with a specific chromosome but has not been assigned an orientation and position. The ‘multi’ category furthest to the right represents NS that are mapping to multiple chromosomes. **B)** Similar results for NS detected in Swe2. **C)** Examples of chromosomal regions where a high amount of NS are detected. The two plots to the left show the localization of 3way overlap sequences (i.e. found in Swe1, Swe2 and HX1) near the centromeric regions of chr14 and chr21. The top right panel displays a region on chr17 where an excess of NS found only in Swe1 and Swe2 could be anchored. The bottom left panel shows NS detected only in the two males (Swe1 and HX1) that could be anchored to regions close to the telomere of chromosome Y.

### Anchoring novel sequences on human chromosomes

We next aimed to anchor the NS onto human chromosomes using information provided by the PacBio long-read data and BioNano optical maps (see Methods section). Only a minority of sequences could be placed into the human genome using this approach, but this analysis still provided valuable insights about the genomic localization of the NS (see **Figure 3B)**. For Swe1 and Swe2, 2.08 Mb and 1.97 Mb of NS, respectively, could be uniquely anchored to a chromosome, while 1.70 Mb and 1.55 Mb were anchored to multiple chromosomes (see **supplementary Table S11**). This again shows that many NS contain repetitive or transposable elements. For the uniquely anchored NS, we observed an accumulation at certain chromosomes. The highest amount of 3-way overlap sequences is present on chromosome 21, while chromosomes 13, 14, and 22 also show enrichment (**Figure 3C**). The NS placed on these chromosomes are mainly localized to centromeric or telomeric regions, suggesting that placement of these sequences has previously been difficult to determine due to their repetitive content. Interestingly, we detected an accumulation of population-specific NS present in both Swedish individuals but not in the Chinese HX1 on chromosome 17. A relatively large amount of sequences (121 kb) shared only between the two male individuals (Swe1 and HX1) could be anchored to the Y chromosome. Surprisingly, we also noted an accumulation of NS shared between Swe1 and HX1 on chromosome 17.

### Application of novel sequences for population scale WGS analysis

Having identified several megabases of DNA not present in the human reference, we were interested to see whether these NS would improve the results of whole genome re-sequencing of the Swedish population. We therefore created a new reference consisting of hg38 combined with all the NS detected in Swe1 and Swe2 (named hg38+NS), after which we leveraged the Illumina WGS data from the SweGen dataset^5^ and aligned the reads from 200 individuals both to hg38 as well as to hg38+NS. The aim of this analysis was to study whether the number of single nucleotide variants (SNVs) was altered as a result of appending NS to the reference, through an analysis procedure outlined in **Figure 4A**.

**Figure 4.**
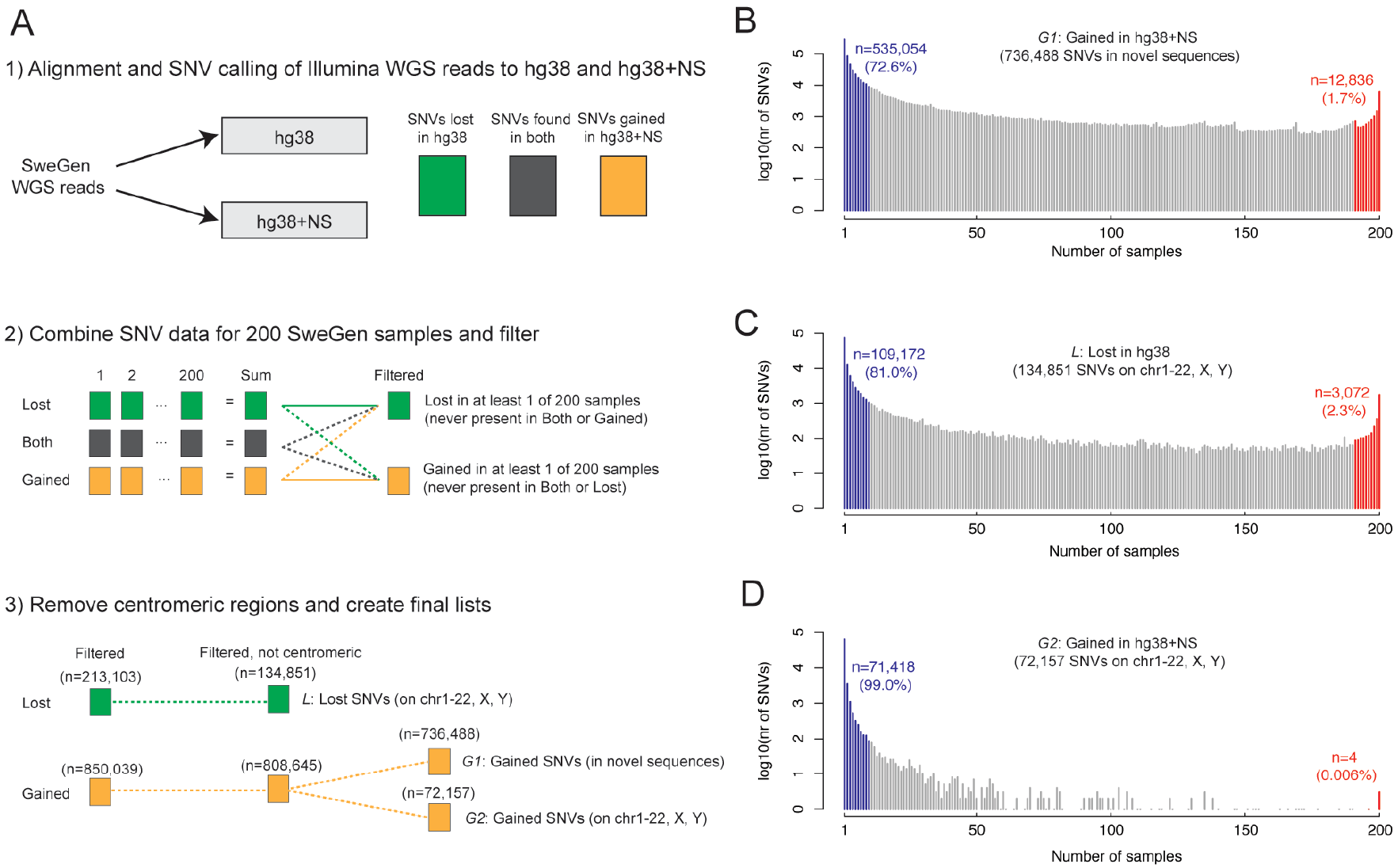
Re-analysis of Illumina WGS data using a Swedish human reference. **A)** Overview of our method to evaluate the effect of NS on SNVs calls from Swedish Illumina WGS data. In the first step, reads from 200 SweGen samples^5^ were aligned both to hg38 and to an extended reference (hg38+NS) where 17.3 Mb of novel sequences detected in Swe1 and Swe2 were appended to hg38. In step 2, SNVs for each of the samples were sorted into three groups: i) SNVs found only in hg38, but not in hg38+NS (named ‘Lost’, in green) ii) SNVs found both in hg38 and hg38+NS (‘Both’, in grey) and iii) SNVs found only in hg38+NS but not in hg38 (‘Gained’, in orange). After such SNV tables were generated for all 200 individuals, a summary file was created for the ‘Lost’ and ‘Gained’ group. The ‘Lost’ SNVs were not allowed to be detected in any of the ‘Gained’ or ‘Both’ files. A similar filtering was performed also for the ‘Gained’ group. In step 3, we further filtered the SNV lists by removing all centromeric regions (from file ‘centromeres_UCSC_hg38.txt’). The resulting ‘Gained’ SNVs were separated into two distinct groups, those present in hg38 chromosomes (chr1-22, X or Y) and those present in the NS. **B)** Frequency distribution of the 736,488 SNVs that were gained in the NS. The x-axis shows the number of SweGen samples (out of 200), and the y-axis shows the number of gained SNVs for each number of samples on a log10-scale. Most of the gained SNVs are detected only in a few samples. The blue and red areas show the number of SNVs that are gained in at most 5% and at least 95% of samples, respectively. **C)** Frequency distribution of the SNVs that were lost in hg38 when adding NS to the hg38 reference. **D)** Frequency distribution of the gained SNVs on chromosomes 1-22, X or Y (i.e. not in novel sequences) when adding NS to the hg38 reference.

An average of 42.5 SNVs per kb was detected in the NS and their frequency distribution is shown in **Figure 4B**. Some of these SNVs are likely to represent true novel genetic variation in the cohort, but a substantial fraction may also originate from errors in the Swe1 and Swe2 assemblies. Surprisingly, the addition of NS to the reference had a large effect on variant calls of the autosomes and sex chromosomes, where 134,851 SNVs disappeared (outside of centromeric regions) when the extended reference was used (**Figure 4C**). These SNVs originate from reads that preferentially aligns to a novel sequence and can be considered as false positives in hg38. Interestingly, we also found a substantial number of SNVs (n=72,157) which were gained on the hg38 chromosomes when using the extended reference. These gained SNVs in hg38 have overall lower allele frequencies compared to the lost SNVs (see **Figure 4D**).

Finally, we investigated SNVs that were consistently lost or gained in hg38 for at least 5% of the 200 SweGen samples when using the extended reference (see **Figure 5A)**. Only a small number of SNVs (n=823) were gained on the hg38 chromosomes in at least 5% of the samples when using the extended reference. However, 26,724 SNVs were lost in at least 5% samples when appending NS to the hg38 reference. These consistently lost SNVs have an uneven distribution over the genome with the highest peak on chrY and smaller peaks on several other chromosomes. Global annotation of the consistently lost SNVs showed that 7,130 (27%) of these are present in version 147 of dbSNP. For the consistently gained SNVs, only 130 (16%) are present in dbSNP, suggesting that these SNVs are more difficult to detect using the hg38 reference alone. A total of 109 consistently lost SNVs were located in a coding sequence of a gene, but none of the consistently gained SNVs were in coding regions. **Figure 5B** shows an example region on chr17 where the NS improved the alignment of Illumina WGS data for two SweGen individuals, resulting in the removal of around 100 false positive SNVs, and importantly, the discovery of seven novel SNVs that were previously masked by the mis-aligned reads. A second example is shown in **Figure 5C** where a region on chrY with about 1000X coverage and many dubious SNVs is cleaned up when NS are appended to hg38. In a third example, as illustrated by the genome browser view of the *FRG2C* locus, the hg38+NS reference improves alignments in coding regions (see **Figure 5D)**.

**Figure 5.**
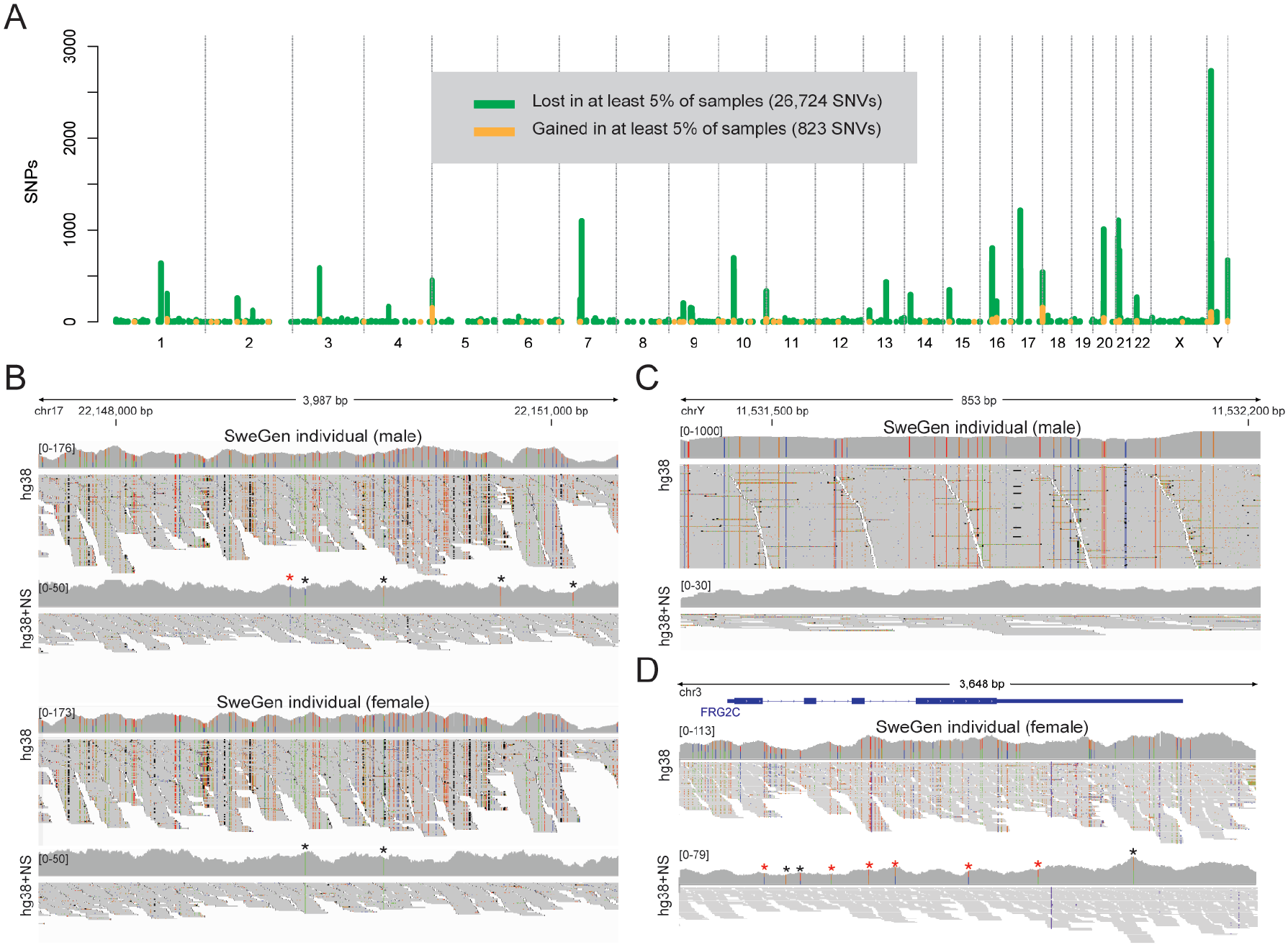
A novel reference gives improved alignment and SNV calling of SweGen WGS data. **A)** Genomic distribution of SNVs that are lost (green) and gained (orange) when NS are appended to the hg38 reference. Only non-centromeric SNVs that are lost/gained in at least 5% of the 200 SweGen samples are shown in this figure. **B)** An IGV^33^ view of Illumina reads for two representative SweGen samples at a region on chr17, where some SNVs are lost and others are gained when using the hg38+NS reference. Illumina data is shown for a male and a female (not the same individuals as Swe1 and Swe2). Both for the male and female, the coverage decreases over the region when NS are appended to hg38, and about 100 (homozygous) false positive SNV calls are lost in each of the samples. Only five heterozygous SNVs where found for the male individual when the novel reference was used, and two homozyogous SNVs for the female (marked by asterisks ‘*’). A red asterisk indicates a gained SNV that is not detected in hg38. **C)** An example region on chrY where the coverage was reduced from almost 1000X to below 30X when using hg38+NS, and where a large number of SNVs were lost. Only data for the male individual is shown in this panel. **D)** Improved alignment and SNV calling over the *FRG2C* locus on chromosome 3. A large number of SNVs were lost, and six SNVs were gained (red asterisks ‘*’) in the female SweGen sample. Some of the lost and gained SNVs are located in the coding sequences of *FRG2C*.

## Discussion

Swe1 and Swe2 represent two of the most complete individual human *de novo* assemblies produced to date. On average, the primary contigs contains 2.87 Gb of unambiguous (non-N) sequence per individual, which is similar to the Chinese HX1^16^ and the Korean AK1^15^ individuals, and 133 million more bases than what could maximally be assembled in any of the 150 Danish genomes^10^. Nevertheless, the Swedish assemblies are still not entirely complete and they could be further improved by BAC sequencing of specific regions^15^, by chromosome interaction mapping^21^, by linked-reads from 10X Genomics^13,15^, or long reads from nanopore-based sequencing^22^. Also, the new technologies make it possible to assemble human genomes starting directly from tissues or blood, instead of cell lines that were used for example for GRCh38. This should resolve the potential issue with genomic aberrations introduced during cell line transformation and long-term culturing^23^.

The Swedish *de novo* assemblies reveal a large amount of NS not present in the GRCh38 reference. Over 5 Mb of NS were overlapping between our two Swedish genomes and the Chinese HX1, and this likely represents valid human DNA sequence. In addition, 1.36 Mb of male-specific sequence was found (i.e. overlapping between Swe1 and HX1) and large fraction of these sequences could be anchored to the Y chromosome. We estimate the total amount of sequence missing from GRCh38 to be at least 6 Mb, which corresponds to about 0.2% of the size of the human genome. Another interesting class of NS are the ~1.5 Mb found in Swe1 and Swe2, but not in HX1. Several of these potentially population-specific NS are clustered at certain genomic regions, such as chr17 (see **Figure 3C**). Some of these regions could have been targets for selection during human evolution, although this needs to be investigated further. In general, the NS are highly repetitive and have elevated GC-contents. At this point, there is no evidence for the presence of functional elements within the NS, although preliminary data suggest that some of them are actively transcribed (data not shown). A more thorough analysis is required to shed light on the functional relevance of these NS.

Our results show that the NS can be used to construct a new version of the human genome reference that improves the analysis of population-scale Illumina WGS data. On average 10,898 SNVs per individual were lost, and 75,035 SNVs per individual were gained in 200 SweGen samples^5^ when appending the NS to GRCh38, with some of this variation affecting the coding sequences of known genes. Because of the stringent filtering options used in our analysis (see **Figure 4A**) these numbers should be seen as a conservative estimate of the novel variation that could be resolved using an improved reference. In addition, since many regions still show poor alignments for SweGen data also when using hg38+NS (data not shown), it is likely that our reference could be further improved and customized for the Swedish population. The benefits of an improved reference are likely to be even stronger for other, non-European, population groups that were poorly represented in the original assembly of GRCh38^9^. In conclusion, despite all efforts to refine the human genome since its original release in 2001^24^, our results indicate that substantial improvements could still be made, not least to represent specific population groups, by *de novo* assembly of representative human genomes from different populations.

## Online methods

### Samples

Swe1 and Swe2 were selected from the 1000 individuals included in the SweGen project^5^. Samples of whole blood from these two individuals were collected in 2006 and frozen without separation of white and red blood cells at −70°C on site, as part of the Northern Sweden Population Health Study (NSPHS) which aims to study the medical consequences of lifestyle and genetics. Genomic DNA was extracted using organic extraction. The NSPHS study was approved by the local ethics committee at the University of Uppsala (Regionala Etikprövningsnämnden, Uppsala, 2005:325 and 2016-03-09). All participants gave their written informed consent to the study including the examination of environmental and genetic causes of disease in compliance with the Declaration of Helsinki.

### PacBio library preparation and sequencing

Four PacBio libraries were produced for each of the Swe1 and Swe2 samples using the SMRTbell™ Template Prep Kit 1.0 according to manufacturer’s instructions. In brief, 10 μg of genomic DNA per library was sheared into 20 kb fragments using the Megaruptor system, followed by an exo VII treatment, DNA damage repair and end-repair before ligation hair-pin adaptors to generate SMRTbell™ libraries for circular consensus sequencing. Libraries were then subjected to exo treatment and PB AMPure bead wash procedures for clean-up before they were size selected with the BluePippin system with a cut-off value of 9500 bp. The libraries were sequenced on the PacBio RSII instrument using C4 chemistry and P6 polymerase and 240 min movie time in a total of 225 SMRTcells™ per sample.

### De novo assembly of SMRT sequencing reads

Raw data was imported into SMRT Analysis software 2.3.0 (PacBio) and filtered for subreads longer than 500 bp or with polymerase read quality above 75. A *de novo* assembly of filtered subreads was generated using FALCON^25^ assembler version 0.4.1 (configuration file is provided as Supplementary Information). In order to improve the accuracy of the assembly, two rounds of sequence polishing were performed using Quiver consensus calling algorithm^25^. For subsequent analysis, primary contigs shorter than 20kb were excluded from the Swe1 and Swe2 assemblies to reduce putative assembly errors. This is slightly more conservative as compared to the Korean AK1 study^15^, where a 10 kb cut-off of primary contigs was used.

### Generation of BioNano optical maps and hybrid assembly

DNA extraction for optical maps was performed at BioNano Genomics (CA, USA), starting from frozen blood from Swe1 and Swe2. Optical mapping was performed on the Irys system (BioNano Genomics) using the two labeling enzymes BssSI and BspQI for each individual. The resulting data was used for a 2-step hybrid assembly of the PacBio contigs using the IrysView software.

### The hg38 reference genome

The hg38 reference genome used in this study is identical to the full analysis set of GRCh38 (accession GCA_000001405.15) described in a study by Zheng-Bradley et al (2017)^18^. This implies that hg38 consists of the primary GRCh38 sequences (autosomes and chromosome X and Y), mitochondria genome, un-localized scaffolds that belong to a chromosome without a definitive location and order, unplaced scaffolds that are in the assembly without a chromosome assignment, the Epstein-Barr virus (EBV) sequence (AJ507799.2), ALT contigs, and the decoy sequences (GCA_000786075.2).

### Quality control and alignment of the two Swedish de novo assemblies

Components of MUMmer3^26^ (NUCmer, delta-filter and dnadiff) were used to assess the quality of the Swe1 and Swe2 *de novo* assemblies and to perform genome alignments. NUCmer (-maxmatch −l 150 −c 400) was used to align each of the assemblies to hg38. After the alignments, delta-filter (-q) was used to filter out repetitive alignments and to keep the best alignment for each assembled contig. Summary statistics for the filtered whole genome alignments were generated by dnadiff.

### Detection of structural variation in PacBio data

We utilized NGMLR (v0.2.3) (https://github.com/philres/ngmlr) and Sniffles^27^ (v1.0.5) to detect structural variations from PacBio long reads. Filtered subreads (min subread length 500 bp and polymerase read quality above 75) were first aligned to the hg38 using NGMLR with default parameters. Sniffles (-s 10 −l50) was subsequently used to identify structural variations >=50bp with at least 10 reads support. Only SV detected on chr1-22, X and Y from Swe1, and on chr1-22 and X from Swe2 were kept for analysis.

### Detection of novel sequences

To identify NS we performed two rounds of sequence mapping using NUCmer^26^. The first round of mapping was the same as described above, when aligning contigs to the hg38 reference. Contigs and part of contigs that failed to align to the reference in the first step were then processed in a second round of mapping, where more relaxed settings were used in an attempt to have more sequences aligned by NUCmer (-maxmatch −l 100 −c 200). Duplicated sequences were then removed to obtain a set of novel sequences (NS) for each of the two indivduals (Swe1 and Swe2). All NS in the final set have a sequence length of at least 100bp with a sequence identity to the hg38 reference that is less than 80%.

### Repeat analysis and BLAST comparison of novel sequences

Repeats in NS were analyzed using RepeatMasker (-species human -s –x; http://www.repeatmasker.org). NS were searched against the nucleotide collection database using BLAST^28^ (2.2.31+) (-max_target_seqs 1 –task blastn –num_threads 16). In order to obtain matched sequences of relatively high similarity, the BLAST results were post processed by setting an E-value threshold at 10^−50^ and by keeping only the top hit for each novel sequence.

### Anchoring novel sequences on human chromosomes

To determine the potential genomic position of the NS that may be anchored, we first used NUCmer (-maxmatch −l 100 −c 200) and delta-filter (-q) to map the NS to the hybrid scaffolds that were generated by PacBio contigs and BioNano optical maps. After anchoring of the hybrid scaffolds, NUCmer (-maxmatch −l 150 −c 400) and delta-filter (-q) were ran to identify the location of the alignments on hg38 chromosomes. NS that mapped to anchored hybrid scaffolds were further analyzed to identify unique or multiple location anchors. NS that were anchored to decoy sequences included in the hg38 reference were excluded from final results.

### Construction of an extended reference based on Swedish novel sequences

Novel sequences detected in Swe1 and Swe2 were appended to hg38 to create an extended version of the human reference sequence (named hg38+NS). For novel sequences overlapping between both Swedish individuals, only the Swe1 version of the NS was used. The resulting hg38+NS reference added 17.3 Mb of novel sequence to hg38.

### Re-alignment of SweGen Illumina data to hg38 and hg38+NS

200 of the SweGen samples^5^ were processed with the Cancer Analysis Workflow (CAW) pipeline (https://github.com/SciLifeLab/CAW) in normal-only mode (no tumor samples), once with hg38 and again with hg38+NS as reference. CAW implements a workflow based on GATK best practices. In summary, reads were aligned using BWA-MEM^29^ 0.7.15with the ALT-aware option turned off. Duplicates were then marked with Picard’s MarkDuplicates 2.0.1. The tools RealignerTargetCreater, IndelRealigner, CreateRecalibrationTable, HaplotypeCaller and GenotypeGVCFs from GATK 3.7.0^30^ were then used in that order to realign around indels, recalibrate base qualities and call variants, respectively, resulting in a final CRAM and VCF file for each sample.

Also, a similar analysis was re-run for 150 of the 200 SweGen samples, but with the ALT aware option turned on in the BWA-MEM alignment. This ALT aware analysis resulted in a much higher number of lost and gained SNVs as compared to the non-ALT aware alignment. We therefore decided to focus on the non-ALT aware analysis in this study, i.e. the analysis run on the 200 samples. By a non-ALT aware alignment we get a conservative estimate of the number of lost and gained SNVs and do not exaggerate the effect of adding novel sequences to the hg38 reference.

### Analysis and annotation of SNVs in SweGen re-alignments

To detect SNVs that were consistently gained or lost among the 200 SweGen samples when the novel sequences were added to hg38, we employed the filtering strategy outlined in **Figure 4A**, implemented by custom scripts in perl and R. ANNOVAR^29^ was used to annotate gained and lost SNVs with information about human genetic variation from dbSNP^30^ v147 and protein coding genes from the NCBI RefSeq database^31^.

## Data Availability

The Swe1 and Swe2 raw sequence data and assembly files will be made available during the spring of 2018 from a local Swedish installation of the European Genome-phenome Archive (EGA) (https://www.ebi.ac.uk/ega) that is now being implemented at Uppsala University and SciLifeLab. The following data files are made available as supplementary material:
**Supplementary Data S1:** SV results for Swe1 and Swe2
**Supplementary Data S2:** Novel sequences in Swe1 and Swe2
**Supplementary Data S3:** BLAST hits in novel sequences
**Supplementary Data S4:** The hg38+NS reference

## Acknowledgements

This work was funded by Science for Life Laboratory (SciLifeLab) as a National Project, supported by the Knut and Alice Wallenberg Foundation (2014.0272), and the Swedish Research Council (PI:UG). PacBio SMRT sequencing was performed by the National Genomics Infrastructure (NGI), which is hosted by SciLifeLab in Uppsala. Optical maps were generated at BioNano (CA, USA). Computations were performed on resources provided by SNIC through Uppsala Multidisciplinary Center for Advanced Computational Science (UPPMAX) under projects b2015225 and sens-2016003. MM and PO were financially supported by the Knut and Alice Wallenberg Foundation as part of the National Bioinformatics Infrastructure Sweden at SciLifeLab.

## References

1. Wong, L.P. et al. Deep whole-genome sequencing of 100 southeast Asian Malays. Am J Hum Genet 92, 52–66 (2013).

2. Boomsma, D.I. et al. The Genome of the Netherlands: design, and project goals. Eur J Hum Genet 22, 221–7 (2014).

3. Gudbjartsson, D.F. et al. Large-scale whole-genome sequencing of the Icelandic population. Nat Genet 47, 435–44 (2015).

4. Fakhro, K.A. et al. The Qatar genome: a population-specific tool for precision medicine in the Middle East. Hum Genome Var 3, 16016 (2016).

5. Ameur, A. et al. SweGen: a whole-genome data resource of genetic variability in a cross-section of the Swedish population. Eur J Hum Genet 25, 1253–1260 (2017).

6. Nakatsuka, N. et al. The promise of discovering population-specific disease-associated genes in South Asia. Nat Genet (2017).

7. Consortium, U.K. et al. The UK10K project identifies rare variants in health and disease. Nature 526, 82–90 (2015).

8. Telenti, A. et al. Deep sequencing of 10,000 human genomes. Proc Natl Acad Sci U S A (2016).

9. Schneider, V.A. et al. Evaluation of GRCh38 and de novo haploid genome assemblies demonstrates the enduring quality of the reference assembly. Genome Res 27, 849–864 (2017).

10. Maretty, L. et al. Sequencing and de novo assembly of 150 genomes from Denmark as a population reference. Nature 548, 87–91 (2017).

11. Ross, M.G. et al. Characterizing and measuring bias in sequence data. Genome Biol 14, R51 (2013).

12. Chaisson, M.J. et al. Resolving the complexity of the human genome using single-molecule sequencing. Nature 517, 608–11 (2015).

13. Mostovoy, Y. et al. A hybrid approach for de novo human genome sequence assembly and phasing. Nat Methods 13, 587–90 (2016).

14. Pendleton, M. et al. Assembly and diploid architecture of an individual human genome via single-molecule technologies. Nat Methods 12, 780–6 (2015).

15. Seo, J.S. et al. De novo assembly and phasing of a Korean human genome. Nature 538, 243–247 (2016).

16. Shi, L. et al. Long-read sequencing and de novo assembly of a Chinese genome. Nat Commun 7, 12065 (2016).

17. Chin, C.S. et al. Phased diploid genome assembly with single-molecule real-time sequencing. Nat Methods 13, 1050–1054 (2016).

18. Zheng-Bradley, X. et al. Alignment of 1000 Genomes Project reads to reference assembly GRCh38. Gigascience 6, 1–8 (2017).

19. Altschul, S.F., Gish, W., Miller, W., Myers, E.W. & Lipman, D.J. Basic local alignment search tool. J Mol Biol 215, 403–10 (1990).

20. Bennett, H.M. et al. The genome of the sparganosis tapeworm Spirometra erinaceieuropaei isolated from the biopsy of a migrating brain lesion. Genome Biol 15, 510 (2014).

21. Bickhart, D.M. et al. Single-molecule sequencing and chromatin conformation capture enable de novo reference assembly of the domestic goat genome. Nat Genet 49, 643–650 (2017).

22. Jain, M. et al. Nanopore sequencing and assembly of a human genome with ultra-long reads. bioRxiv (2017).

23. Redon, R. et al. Global variation in copy number in the human genome. Nature 444, 444–54 (2006).

24. Lander, E.S. et al. Initial sequencing and analysis of the human genome. Nature 409, 860–921 (2001).

25. Chin, C.S. et al. Nonhybrid, finished microbial genome assemblies from long-read SMRT sequencing data. Nat Methods 10, 563–9 (2013).

26. Kurtz, S. et al. Versatile and open software for comparing large genomes. Genome Biol 5, R12 (2004).

27. Sedlazeck, F.J. et al. Accurate detection of complex structural variations using single molecule sequencing. bioRxiv (2017).

28. Camacho, C. et al. BLAST+: architecture and applications. BMC Bioinformatics 10, 421 (2009).

29. Wang, K., Li, M. & Hakonarson, H. ANNOVAR: functional annotation of genetic variants from high-throughput sequencing data. Nucleic Acids Res 38, e164 (2010).

30. Sherry, S.T. et al. dbSNP: the NCBI database of genetic variation. Nucleic Acids Res 29, 308–11 (2001).

31. O’Leary, N.A. et al. Reference sequence (RefSeq) database at NCBI: current status, taxonomic expansion, and functional annotation. Nucleic Acids Res 44, D733–45 (2016).

32. Genomes Project, C. et al. A global reference for human genetic variation. Nature 526, 68–74 (2015).

33. Thorvaldsdottir, H., Robinson, J.T. & Mesirov, J.P. Integrative Genomics Viewer (IGV): high-performance genomics data visualization and exploration. Brief Bioinform 14, 178–92 (2013).

